# Utility of brain-derived extracellular vesicles from human umbilical cord blood to measure non-infectious neuroinflammation and functional iron deficiency

**DOI:** 10.1101/2025.04.15.648967

**Authors:** Phu V. Tran, Shirelle X. Liu, Andrew C. Harris, Zia L. Maxim, Manjula Munirathinam, Peter Muelken, Narmin Javadova, Sowmya Kruttiventi, Pamela J. Kling, Michelle L. Baack, Michael K. Georgieff

## Abstract

**Background:** Non-infectious neuroinflammation (NINI) in early life and neonatal neural iron deficiency (nID) have been proposed to contribute to neurodevelopmental dysfunction and disorders, including autism, through toxic effects on neural cells and impaired molecular signaling. Early detection of NINI and nID may enable interventions to restore neurodevelopmental homeostasis and reduce long-term impact. However, such diagnoses are impossible due to ethical reason to access to the central nervous system (e.g., lumbar puncture) when there is no suspicion of brain infection (e.g., meningitis).

**Methods:** We refined a methodology to isolate and quantify inflammatory and iron-regulatory proteins in high-quality brain-derived extracellular vesicles (BDEVs) from human umbilical cord blood. A preclinical model was used to validate that BDEV contents reflect the brain microenvironment. We then applied this approach to evaluate biomarkers of NINI and nID in BDEVs isolated from cord plasma of newborns of mothers with obesity, a non-infectious pro-inflammatory gestational condition and a risk factor for nID.

**Findings:** Plasma BDEV analytes correlated more strongly with brain analytes and showed stronger associations among functionally related molecules compared with whole plasma analytes. Maternal obesity induced an anti-inflammatory response in the brain compartment, a pro-inflammatory response in the periphery, and functional nID.

**Interpretation:** BDEVs may provide a more sensitive representation of the brain microenvironment than blood-based measures (e.g., plasma), enabling non-invasive, early detection of infants with NINI and neonatal nID.

**Funding:** Supported by NIH, the Masonic Institute for Developing Brain, Hennepin Healthcare Research Institute, UnityPoint Health Meriter Foundation, and American Academy of Pediatrics Resident Research Grant.

**Research in Context:** *Evidence before this study:* Previous studies hypothesized that gestational inflammatory environments (e.g., maternal obesity) increase risk of neonatal neural iron deficiency (nID), neuroinflammation, and neurodevelopmental disorders. However, assessing non-infectious neuroinflammation (NINI) in infants from these high-risk groups is not feasible or clinically justifiable using invasive methods (e.g., cerebrospinal fluid collection), and analytes from whole blood or plasma may not accurately reflect brain physiological status.

*Added value of this study:* We refined a protocol to isolate brain-derived extracellular vesicles (BDEVs) from a small amount of plasma sample (100 µL) and characterized their contents to approximate brain physiology. We demonstrated that BDEVs more accurately reflect brain status compared with whole plasma. In newborns of mothers with obesity during pregnancy, we found evidence of a low-grade peripheral inflammation accompanied by a protective anti-inflammatory response in the brain. Maternal obesity was also found to induce neonatal nID.

*Implications of all the available evidence:* BDEVs can serve as an accurate, non-invasive tool to assess brain condition. In clinical settings, this approach may be used for early diagnosis, monitoring disease progression, and evaluating treatment effects.

**Graphical Abstract:** 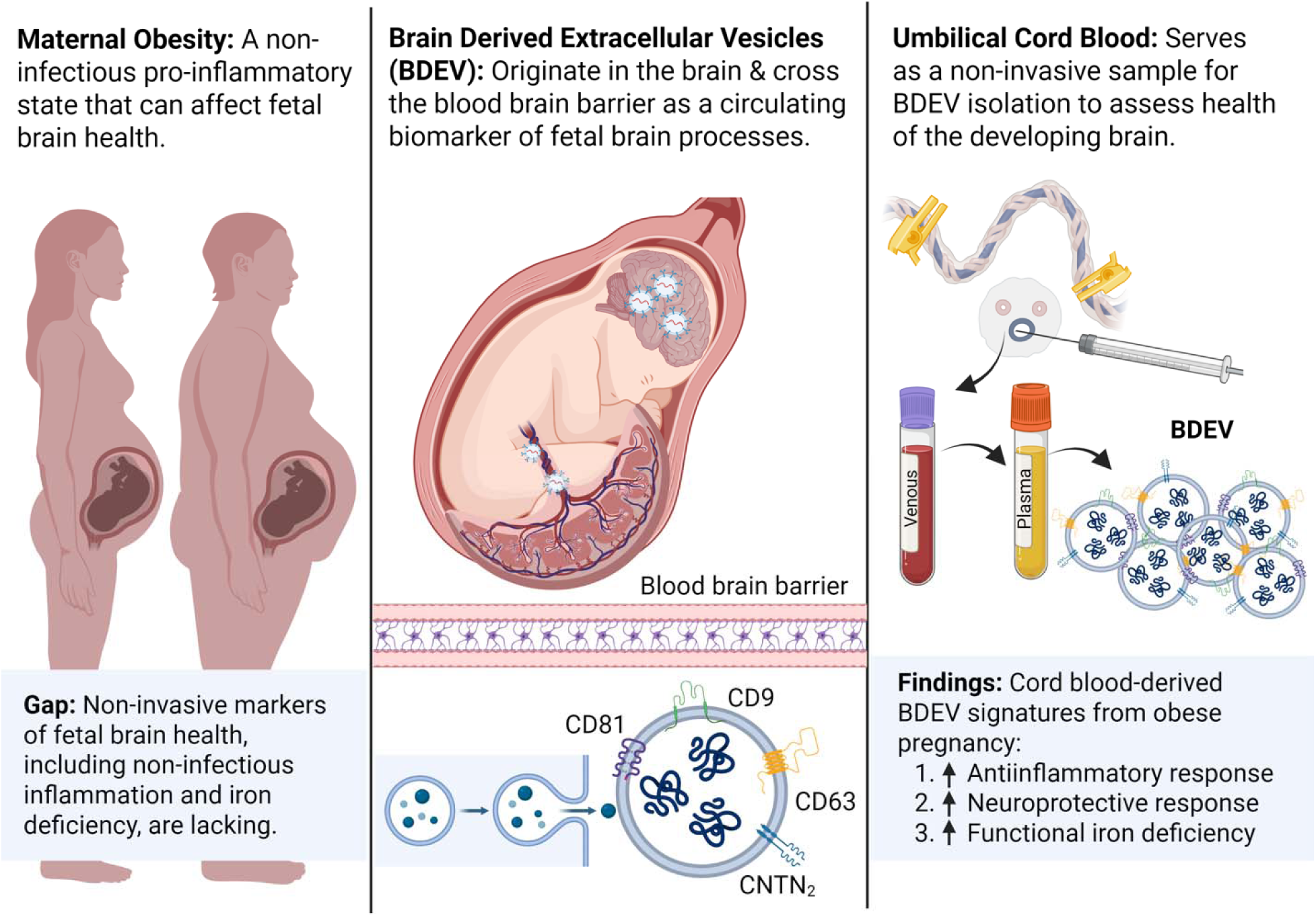

## Introduction

Neurodevelopmental dysfunction represents a broad range of conditions that are associated with multi-faceted care and resource utilization throughout childhood and potentially into adulthood. Non-infectious neuroinflammation (NINI) may occur early in life, particularly in offspring born to women with gestational conditions characterized by an inflammatory state, such as obesity. NINI may contribute to neurodevelopmental dysfunction via the inflammatory cascade’s toxic effects on both neurons and astroglia. Evidence from preclinical models and postmortem human tissue indicate that altered chemokine/cytokine signaling can lead to dysregulated cellular homeostasis (e.g., oxidative stress), reduced cell survival, impaired neuronal differentiation (e.g., myelination, gliosis) and neural iron deficiency (nID) (1–4). These molecular and cellular changes are potential mechanisms underlying the link between NINI and several neurodevelopment disorders, including autism spectrum disorders (ASD), attention deficit hyperactivity disorder (ADHD), and intellectual disability (5–9).

NINI falls within the spectrum of Developmental Origins of Health and Disease (DOHaD), where early-life environment plays a major determining factor for physical and mental health across the lifespan (10). Thus, detection of NINI in early life may provide opportunities for intervention during or after neurodevelopmental events to minimize their long-term impact. Pediatric blood sampling is feasible in clinical practice; however, peripheral blood or serum concentrations of analytes indexing inflammation may not accurately reflect inflammation within the brain. This is because of the contribution of other organs (e.g., liver, spleen) to that pool (11, 12). Because of barriers related to accessing the central nervous system (CNS) compartment non-invasively, there are limited approaches to assessing CNS state in a developing brain, specifically in screening pediatric patients for NINI when cerebrospinal fluid collection is neither indicated, nor ethically feasible.

Currently, there are experimental approaches to develop better CNS screening methods using brain-derived extracellular vesicles (BDEVs), a class of extracellular vesicles that reflect origin cell status and cell-cell communication via the contained cargo of proteins, carbohydrates, lipids, metabolites, and nucleic acids (13, 14). BDEVs cross the blood brain barrier into systemic circulation and have been explored as potential carriers of biomarkers that approximate the current physiological state of brain cells because of their short half-lives which range from minutes to hours (15, 16).

The value of BDEVs in detecting NINI as a non-invasive method is dependent on identification and validation of NINI biomarker signatures. In diagnostic development, clinical biomarkers are often first identified using samples from patient cohorts that have been exposed to risk factors. For NINI, numerous gestational exposures reportedly instigate metabolic derangements and a non-infectious proinflammatory state during pregnancy to perturb neurodevelopment in the developing offspring (17–19). These include excess production of cortisol during chronic stress, exposure to toxins, nutritional deficits, obesity and excessive gestational weight gain (20–22). Specifically, the status of maternal inflammation plays a critical role in programming the fetal brain immune responses, which regulate normal fetal brain development and neuronal maintenance throughout life (20). Emerging evidence recognizes maternal obesity as a risk factor for abnormal programming of the immune system and is associated with peripheral inflammation, nID, and a higher risk of atypical neurodevelopment in offspring (22, 23). Mechanistically, the effects of maternal obesity on neurodevelopment in pre-clinical models have been ascribed to excess neuroinflammatory activation (24, 25), brain iron dysregulation (26–28), and abnormal synaptic connectivity and neurotransmission (29, 30). The results from these preclinical models underscore the translation of NINI and nID assessment of potential clinical biomarkers.

We have previously suggested that BDEVs isolated from human umbilical cord samples had utility to assess neural markers contactin-2 (CNTN2) and brain derived neurotrophic factor (BDNF) (31). Here we describe refinement of the BDEV isolation methodology from human umbilical cord samples and quantify inflammatory and iron regulatory proteins therein. We first utilized a preclinical model to establish that BDEV contents reflect brain tissue status. We then investigated the feasibility of using circulating BDEVs to compare NINI and nID biomarkers in high-and low-risk asymptomatic newborns by leveraging easily accessible umbilical cord blood that is sampled immediately after birth, before the onset of postnatal confounders. The findings indicate that contents within BDEVs may be used to index brain NINI and nID in response to the *in utero* proinflammatory state of maternal obesity. Taken together, BDEVs isolated from umbilical cord blood may serve as an excellent indicator of concurrent maternal-fetal physiological status, including inflammation and iron trafficking across the placental-fetal unit. Moreover, analysis of risks for atypical neurodevelopment early in life provides opportunities within a critical developmental window for close follow-up and timely postnatal interventions that can have maximal impact on development.

## Methods

### Preclinical model to demonstrate BDEV contents as index of brain concentrations

A preclinical rat model was used to demonstrate that BDEVs accurately reflect brain concentrations of inflammatory analytes. All procedures were approved by the Hennepin Health Research Institute IACUC (protocol #22-02). Rat (Sprague-Dawley) pups used in this study were generated by in-house breeding. Dams were kept in a conventional rodent housing facility with ad libitum access to food and water, 22°C room temperature, and 12h/12h light/dark cycle. Postnatal day (P)7, equivalent to human brain development in the 3^rd^ trimester (32), iron-sufficient rat pups (n=10, 1-2 pups/sex from 4 litters) were euthanized by decapitation under isoflurane anesthesia. Trunk blood samples were collected into EDTA (50 µL, 0.5M) solution and plasma was isolated by centrifugation (13,000 RPM, 10 min, room temperature, tabletop centrifuge). Plasma was separated and stored at −80°C. Hippocampus was chosen for this study since it is one of the most metabolically active brain regions that is rapidly developing at this age (33, 34), and thus at greatest risk for NINI and nID. Hippocampi were micro-dissected, flash-frozen in liquid nitrogen, and stored at −80°C until use.

Frozen hippocampus was homogenized in 1.0 mL of cold PBS, centrifuged at 10,000 RPM (5 min), and supernatant was transferred into a new tube. 100 µL of hippocampal homogenate was digested with proteinase K (1 µg/mL, 37°C, 10 min) to remove protein aggregates. Following proteinase K inactivation (on ice, 5 min), small EV were isolated from the digested hippocampal homogenate and paired plasma for analyses as described below.

### Human subjects and biospecimen collection

Bio-banked umbilical cord plasma samples used for this study were initially collected between November 2017 and February 2024 following routine cesarean section delivery of consenting pregnant participants who were enrolled in Sanford Health’s Cord Blood Feasibility study (IRB STUDY00000571: Cord Blood/Tissue Feasibility and Placenta Research). The study was carried out under guidance from Sanford’s Institutional Review Board in accordance with The Federal Regulatory Guidelines (Federal Register Vol. 46, No. 17, January 27, 1891, Part 56) and guidance from the Office of Human Research Protection (45 CFR 46) as described in the 21 CFR 50, Protection of Human Subjects. Pregnancy data were collected from electronic medical records in accordance with U.S. federal Health Insurance Portability and Accountability Act (HIPAA) assurances. Consenting donors between the ages of 18-45 who were pregnant and scheduled for a planned C-section delivery at Sanford Health were included in the study. Subjects with known HIV/AIDs, Hepatitis B, Hepatitis C or other blood borne infectious diseases, those whose delivering physician felt collection of the cord blood, tissue or placenta would affect the health of the mother or infant, or those who, in the judgment of the investigator, were unable to give an informed consent for reasons of incapacity, immaturity, adverse personal circumstances or lack of autonomy were excluded from the study.

Overall, the Cord Blood Feasibility Study enrolled 300 maternal donors. Demographic data collected included maternal age, gravidity, parity, pre-pregnancy and post-pregnancy height, weight and BMI. When there was no recent pre-pregnancy weight recorded, a first-visit, first-trimester weight was substituted. Additional data collected included smoking status, pregnancy complications (including preeclampsia and diabetes), newborn gestational age, birth weight, length, head circumference, and sex. This study used only a subset of samples from healthy, women with a range of BMI and singleton pregnancies and no diagnosis of chorioamnionitis, preeclampsia, diabetic pregnancy (Type 1, 2 or gestational), multiple fetal anomalies or abnormal genetic testing.

For group comparisons, pre-pregnancy (or first-visit) BMI was used to cluster participants in underweight (BMI<18.5), normal weight (BMI=18.5-24.9), overweight (BMI=25-29.9) and obese (BMI≥30) categories according to the Institute of Medicine (IOM) and American College of Obstetricians and Gynecologists (ACOG) Clinical Guideline number 548 (Table 1a) (35). Women from this subset who were underweight (n=1) or normal weight (n=21) were clustered with the control group. Women who were overweight (n=14) or obese (n=24) were clustered with the overweight-obese (OWO) group. Immediately post-delivery, the umbilical cord was clamped and cut. Before delivery of the placenta, whole blood was collected by gravity from the cut end of the cord using aseptic technique into a sterile anti-coagulant coated sterile blood collection bag. Cord blood was refrigerated and processed by centrifugation for plasma collection as soon as feasible and always within 12 hours. Plasma was aliquoted, frozen and banked at −80°C until use.

**Table 1.** Demographic data of the studied cohorts.

| Weight Category | Pre-pregnancy BMI | Inadequate GWG | Appropriate GWG | Excessive GWG |
| --- | --- | --- | --- | --- |
| **Underweight** | <18.5 | <12.7 kg<br>(<28 lbs) | 12.7-18.2 kg<br>(28-40 lbs) | >18.2 kg<br>(>40 lbs) |
| **Normal weight** | 18.5-24.9 | <11.4 kg<br>(<25 lbs) | 11.4-15.9 kg<br>(25-35 lbs) | >15.9 kg<br>(>35 lbs) |
| **Overweight** | 25-29.9 | <6.8 kg<br>(<15 lbs) | 6.8-11.4 kg<br>(15-25 lbs) | >11.4 kg<br>(>25 lbs) |
| **Obese** | ≥30 | <5 kg<br>(<11 lbs) | 5-9.1 kg<br>(11-20 lbs) | >9.1 kg<br>(>11 lbs) |

| Demographic | Control (n=22) | OWO (n=38) | p-value |
| --- | --- | --- | --- |
| Pre-pregnancy weight (kg) | 59.74 ± 6.20 | 90.40 ± 19.90 | **<0.0001** |
| Pre-pregnancy BMI | 22.23 ± 1.97 | 33.02 ± 6.75 | **<0.0001** |
| Post-pregnancy weight (kg) | 72.00 ± 7.65 | 99.82 ± 16.19 | **<0.0001** |
| Gestational Weight Gain (kg) | 12.20 ± 5.53 | 9.67 ± 7.45 | 0.18 |
| Inadequate Weight Gain (%) | 41% | 23% | 0.15<sup>a</sup> |
| Appropriate Weight Gain (%) | 32% | 22% | 0.54<sup>a</sup> |
| Excessive Weight Gain (%) | 27% | 54% | 0.06<sup>a</sup> |
| Maternal Age (yr) | 31.82 ± 4.56 | 29.40 ± 4.19 | **0.03** |
| Gravidity (n) | 2.73 ± 1.28 | 3.22 ± 1.25 | 0.15 |
| Parity (n) | 2.23 ± 0.97 | 2.44 ± 1.03 | 0.45 |
| Smoking (%) | 9.10% | 5% | 0.61<sup>a</sup> |
| GA (weeks) | 38.70 ± 0.84 | 38.60 ± 0.97 | 0.62 |
| Birth Weight (kg) | 3.34 ± 0.46 | 3.51 ± 0.46 | 0.22 |
| Birth Weight Percentile | 51 ± 31 | 63 ± 25 | 0.10 |
| Birth Weight Z-score (n) | 0.07 ± 1.01 | 0.45 ± 0.83 | 0.11 |
| Male (%) | 59% | 53% | 0.79<sup>a</sup> |

| Demographic | Inadequate (n=19) | Appropriate (n=15) | Excessive (n=26) | p-value |
| --- | --- | --- | --- | --- |
| Pre-pregnancy weight (kg) | 82.70 ± 27.40 | 77.91 ± 20.47 | 77.75 ± 18.17 | 0.73 |
| Pre-pregnancy BMI | 31.16 ± 10.21 | 28.74 ± 6.92 | 28.18 ± 5.83 | **0.04** |
| Post-pregnancy weight (kg) | 87.32 ± 22.75 | 88.26 ± 17.91 | 93.55 ± 16.89 | 0.49 |
| Gestational Weight Gain (kg) | 4.42 ± 5.91 | 10.39 ± 4.10 | 15.86 ± 4.28 | **<0.0001** |
| OWO Group (%) | 50% | 56% | 79% | 0.09 |
| Age (yr) | 30.72 ± 5.42 | 32.63 ± 4.56 | 28.89 ± 3.52 | **0.03** |
| Gravidity (n) | 3.11 ± 1.23 | 3.12 ± 1.36 | 2.86 ± 1.18 | 0.71 |
| Parity (n) | 2.56 ± 0.92 | 2.13 ± 0.62 | 2.29 ± 1.12 | 0.41 |
| Smoking (%) | 11% | 0% | 7% | 0.47 |
| Preeclampsia (%) | 0% | 6% | 7% | 0.61 |
| GA (weeks) | 38.75 ± 0.72 | 38.59 ± 1.06 | 38.97 ± 1.06 | 0.45 |
| Birth Weight (kg) | 3.33 ± 0.58 | 3.46 ± 0.39 | 3.47 ± 0.42 | 0.54 |
| Birth Weight Percentile | 50.78 ± 18.25 | 62.31 ± 23.76 | 61.54 ± 25.87 | 0.37 |
| Birth Weight Z-score (n) | 0.08 ± 1.11 | 0.47 ± 0.87 | 0.37 ± 0.79 | 0.42 |
| Male (%) | 50% | 69% | 46% | 0.42 |

Data include mean ± standard deviation. Group differences were calculated using one-way ANOVA or \*Fisher's exact test. Significant p values of <0.05 are bold.

### Isolation of small Extracellular Vesicles (sEVs)

Small extracellular vesicles (sEVs) were isolated using the size-exclusion gravity-flow Exo-Spin 96 kit following the manufacturer’s protocol (Cell Guidance Systems, St. Louis, MO, USA) (36). 100 µL hippocampal homogenate (rat) or plasma (rat and human) was applied to each column and allowed to drain by gravity. Flow-through were discarded and excess droplets removed from the nozzle before placing it on top of a hydrophobic collection plate. sEVs were eluted from columns using 100 µL (2x) PBS (137 mM NaCl, 2.7 mM KCl, 8 mM Na_2_HPO_4_, 2 mM KH_2_PO_4_, Cytiva, Logan, UT, USA). The eluted EVs fractions were combined and transferred to clean, labeled 1.5 mL tubes (Supplemental Figure 1a).

### Enrichment of brain-derived extracellular vesicles (BDEVs)

Brain-derived extracellular vesicles (BDEVs) were enriched from sEVs preparation by immunoprecipitation (IP) using antibodies targeting CNTN2 (Cat. # PA5-101541, ThermoFisher Scientific, Carlsbad, CA, USA) (31) (Supplemental Figure 1a). Specifically, isolated plasma sEVs were incubated with 1.5 µg polyclonal rabbit anti-CNTN2 antibody in PBS with end-over-end rotation at 4°C overnight. Antibody concentration was based on optimization by titration assays assessing IP efficacy with 0.5, 1.0, 1.5, and 2.0 µg. 100 µL protein-A Agarose beads (50% slurry, Sigma, St. Louis, MO, USA) were added into a new 1.5 microfuge tube and centrifuged for 1 min (10,000 rpm) and the liquid was removed. sEV and antibody reaction was transferred into the tube containing protein-A agarose beads and mixed for 2 hours, centrifuged 1 min at maximum speed. The supernatant (CNTN2-depleted fraction, CNTN2^dep^) was removed and stored in a new 1.5 mL tube. The sEV/antibody/bead complexes were rinsed (3x) with 500 µL IP wash buffer (25mM Tris-HCl pH 7.4, 150mM NaCl, 1mM EDTA, 1% NP-40, 5% glycerol), centrifuged (3 min, 2500 x g), and supernatant was discarded. BDEVs were eluted twice, each with 100 µL low pH elution buffer (IgG Elution Buffer, Cat. # 21004, ThermoFisher Scientific) and a 5-minute incubation with mixing before centrifugation (2 min, 2500 x g). 10 µL 1M TRIS pH 9.0 was added to neutralize the eluent (Supplemental Figure 1a).

### Quality assessment of BDEVs by nanoparticle tracking analyzer (NTA)

The isolated BDEVs were assessed for concentrations and sizes using a nanoparticle tracking analyzer (ZetaView MONO, 520nm laser, Particle Matrix, Ammersee, Germany). 10 µL BDEV suspension was diluted in 990 µL milli-Q water(1:100), injected into the flow-cell using a 1 mL syringe (Norm-Ject-F Luer Solo, Air-Tite Products Inc, Virginia Beach, VA, USA). Particles were imaged with 11 frames using a dynamic range between 40–400 particles per frame (Software version 8.06.01 SP1) on the EV_max_ SOP (analysis parameters: Max Area 1000, Min Area 10, Min Bright. 18, nm/Class 5, TL 15). Concentration and distribution of particle diameters were compiled from all images and reported as average counts per frame (Supplemental Figure 1b).

### Validation of BDEVs by Transmission Electron Microscopy (TEM)

Three random BDEVs samples from human cord plasma were imaged using TEM to examine their morphology. In brief, BDEV isolates (100 µL) were delivered to the Characterization Facility at the University of Minnesota for TEM imaging. TEM images were captured by negative staining samples with 1% Uranyl Acetate (UA) on glow discharged 400 mesh carbon coated Cu grids and imaging with FEI Tecnai Spirit Bio-Twin TEM at 120kV (Hillsboro, OR, USA). Images were captured at 9,300x, 13,000x, and 18,500x magnifications (Supplemental Figure 1c).

### Validation of BDEVs by dot blots and ELISA

Validation of BDEVs was performed as previously described (31, 37). In brief, 5 µL of BDEVs were assayed for sEV markers (CD9, CD63, CD81, mouse monoclonal, Invitrogen Inc., Carlsbad, CA, USA), biogenesis markers (TSG101, Calnexin, mouse monoclonal, Invitrogen Inc.), and neural-specific markers (NF-L, mouse monoclonal, Abcam, Waltham, MA, USA, and NMDAR2A, mouse monoclonal, Invitrogen Inc.) by dot blots (see Supplemental Table 1 for Cat. #). Paired samples of plasma, total sEVs, CNTN2-depleted sEVs, and CNTN2-positive sEVs were dotted onto nitrocellulose paper. Blots were blocked in 6 mL of blocking buffer for near IR fluorescent imaging (Rockland, Pottstown, PA, USA) for 1 hour. Following the removal of blocking solution, blots were incubated in primary antibody (1:10,000 diluted in blocking buffer) at 4°C overnight. Blots were then washed 3x with PBS + 0.1% Tween20 (PBST) and incubated in fluorescent-conjugated secondary antibody (AF680, goat anti-mouse IgG, 1:12,500 dilution) in blocking buffer for 45 minutes at RT. Blots were washed with PBST (3x) and PBS, then scanned using a Li-Cor Odyssey XF instrument (Licor-bio, Lincoln, NE, USA) (Supplemental Figure 1d). To further validate BDEVs, random samples were assayed using the Human Exosome Characterization Panel that assesses the expression of CD9, CD63, CD81, Syn-1, Flot-1, TSG101, Ago2, CALR, and GAPDH (MilliporeSigma, HEXSM-170-PMX) per MISEV2023 guidelines (38) (Supplemental Figure 1e) and the Luminex platform (see below).

### Assays for quantifiable cytokines, chemokines, and iron protein markers associated with BDEVs and paired hippocampal tissue exosomes of P7 rat pups

Samples were assessed by the Cytokine Reference Laboratory (CRL, University of Minnesota) using commercially available kit/reagents. This is a CLIA’88 licensed facility (license #24D0931212). Rat hippocampal sEV (10^9^ particles) and paired plasma BDEVs (10^9^ particles) were analyzed using the rat cytokine panel, which consists of 27-analytes (Milliplex, Millipore, St. Louis, MO, USA). Following the preparation of all reagents, samples, and standards, 50 µL of standard or sample was added to each well in duplicates. 50 µL of diluted microparticle cocktail (fluorescent color-coded beads coated with a specific capture antibody) was added to each well. The samples were incubated for 2 hours at room temperature (RT) on a shaker at 800 rpm. After removing the liquid from each well, reactions were rinsed with 100 µL Wash buffer (3x). 50 µL diluted Biotin-Antibody cocktail was then added to each well and incubated for 1 hour at RT on the shaker (800 rpm). The samples were then rinsed with 100 µL wash buffer (3x), replaced with 50 µL diluted Strepavidin-PE, and incubated for 30 min at RT with shaking (800 rpm). Following the removal of liquid, 100 µL Wash buffer was added to the samples and signals were read using the xMAP INTELLIFLEX (R&D Systems, Minneapolis, MN, USA) instrument equipped with a dual-spectra laser, which detects the color-coded beads (analytes) and PE signals (concentration of analytes). Concentration of analytes was determined by extrapolation from 4- or 5-parameter fitted standard curves using Belysa Immunoassay Curve-Fitting software (R&D Systems).

### Quantification of analytes (chemokines, cytokines, iron protein markers) in human BDEVs and plasma

For BDEVs, 1×10^9^ particles from human umbilical cord plasma samples were assayed using a customized panel with Fractalkine (CX3CL1), TNFα, Eotaxin (CCL11), CRP, BDNF, PARK7, S100B, Ferritin, and TfR (R&D Systems) using the Luminex platform (xMAP INTELLIFLEX) and magnetic bead set (Cat# LXASHM-09) following manufacturer’s protocol (R&D Systems) as described above. Selection of analytes were based on a literature-reviewed proposed working model for interactions among neural cell types at molecular levels with a specific focus on immune response and associated cellular iron homeostasis (Fig. 1).

**Fig. 1:**
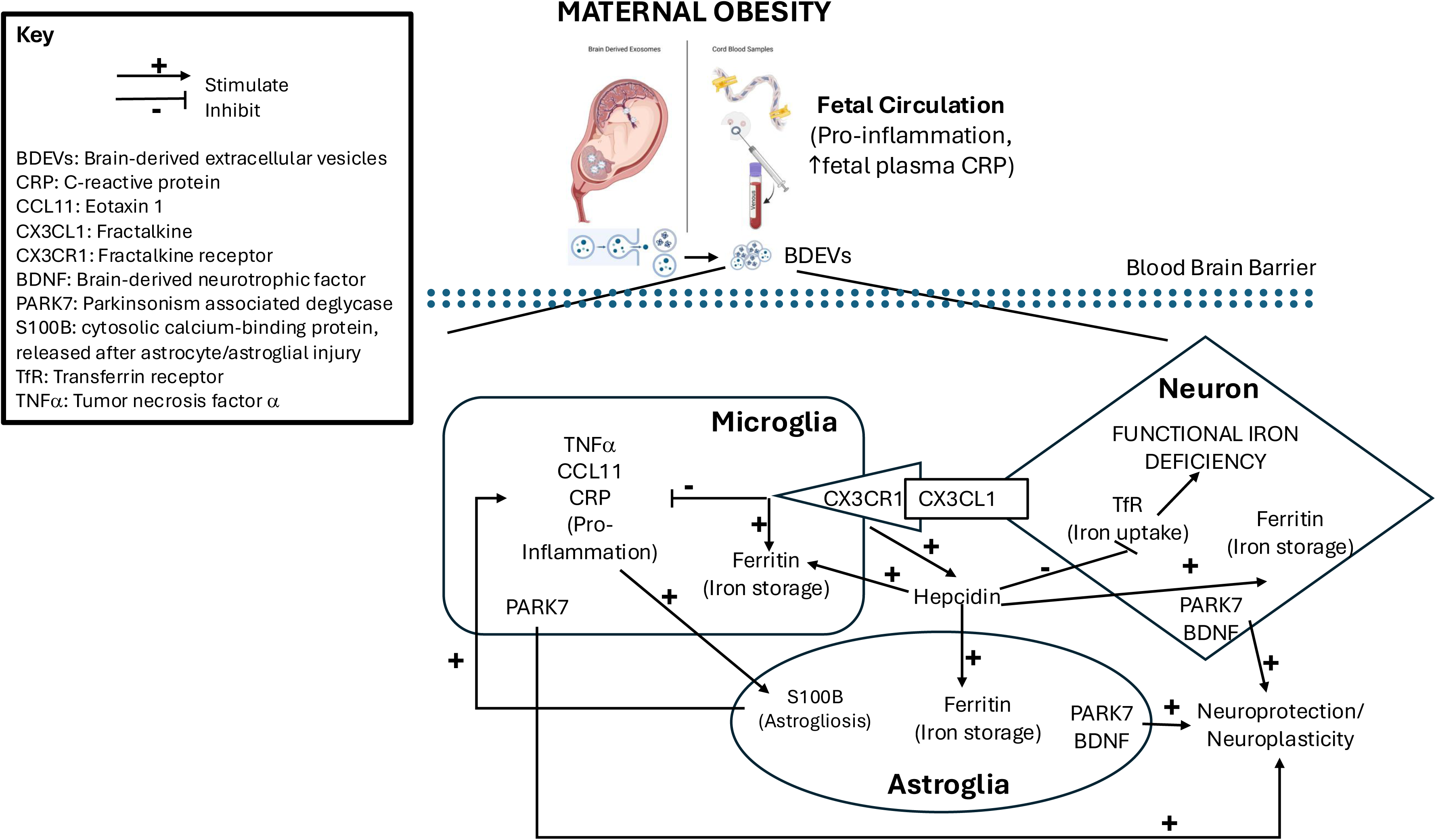
A proposed model of brain iron indices and immune responses in fetus/newborn of mother with obesity through altered maternal-fetal metabolic milieu on microglia, astrocytes, and neurons, including the neuron-microglia CX3CL1/CX3CR1 signaling and astrocyte-microglia S100B signaling.

For plasma analysis, an aliquot (10 µL) of umbilical cord plasma was assayed for concentrations of the same analytes in the above panel. The quantification was performed by the CLIA’88 Cytokine Lab using the Luminex platform as described above.

### Quantification of hepcidin in plasma and BDEVs

For plasma hepcidin quantification, 10 µL plasma samples (n=20/group) was assayed using single plex ELISA following manufacturer’s protocol (R&D Systems).

To generate a fluorescent unit (FU) to protein concentration (pg/mL) conversion standard curve, the same samples were used in a dot array assay. This was done in order to approximate the concentration of BDEVs using the dot array method as they were below the detection limit of the ELISA method. In brief, 4 µL diluted plasma (1:10 dilution in PBS) were dotted in duplicates onto a nitrocellulose membrane using a 96-well grid. Blot was blocked in 6 mL of blocking buffer for near IR fluorescent imaging (Rockland) for 1 hour and then replaced with anti-hepcidin antibody (rabbit polyclonal, 1:20,000 dilution, R&D Systems) in blocking buffer and incubated at 4°C overnight. Blot was washed with PBST (3x) and incubated in fluorescent-conjugated secondary antibody (A700, goat anti-rabbit IgG, 1:12,500 dilution) in blocking buffer for 45 minutes at RT, washed with PBST (3x), and scanned using a Li-Cor Odyssey XF (Licor-bio, Lincoln, NE, USA) instrument. The results generated a FU-protein concentration (pg/mL) conversion curve: *y*(*FU*) = 0.027 * *x*(*ng*/*ml*) + 0.0549 with a linear regression coefficient of determination R^2^ = 0.95, p = 0.02.

For BDEV hepcidin quantification, dot array was performed. In brief, 4 µL of BDEV isolates were dotted in duplicates onto a nitrocellulose membrane using a 96-well grid. BDEV concentrations (particles/mL) were determined using the nanoparticle analyzer (ZetaView). Blot was processed as described above with an additional blocking step with goat anti-rabbit IgG (1:2000 diluted in blocking buffer) to quench any residual rabbit IgG in BDEVs prior to incubation with rabbit anti-human hepcidin antibody (1:20,000 dilution in blocking buffer). Fluorescent intensity unit (FU) for individual dot was quantified using Image Studio software (v5.2, Licor-bio) and normalized to the total volume (4 µL) and transposed to pg/10^9^ particles using a previously determined ELISA-based measurement unit conversion standard curve above.

### Statistical methods

Prism (GraphPad v10.5.0, Boston, MA, USA) was used for statistical analyses. Data were processed using 0.1% ROUT method to identify outliers. Spearman r correlation matrix was generated for continuous variables of plasma and BDEV data. The correlation matrices were graphed using MS Excel with significant values highlighted. Non-linear regression analyses were used to determine the correlation between variables and groups. Group comparisons were made by Welch’s *t*-test or non-parametric Mann-Whitney U-test to account for unequal variances (OWO vs. Controls) for continuous variables, Fisher’s exact test for categorical variables (e.g., smoker vs non-smoker, newborn sex), and Brown-Forsythe and Welch ANOVA tests to account for unequal standard deviation. Significance was defined as p<0.05. This study was not powered to consider newborn sex as a biological variable. Cord blood samples from both sexes were included in all analyses.

## Results

### Correlations between brain sEVs and CNTN2^+^ sEVs in rats

To demonstrate that BDEVs accurately reflect brain concentrations of the analytes, a preclinical experiment was designed to compare the correlation between CNTN2^+^ sEVs extracted from plasma and sEVs extracted directly from hippocampus of postnatal day 7 rats (Fig. 2a). Assessment of paired plasma CNTN2^+^, CNTN2^dep^, and hippocampal sEVs revealed that CNTN2^+^ sEVs, but not CNTN2^dep^ sEVs, positively correlated with hippocampal sEV for TNFα and IL-2, two quantifiable cytokines (Figs. 2b-c).

**Fig. 2:**
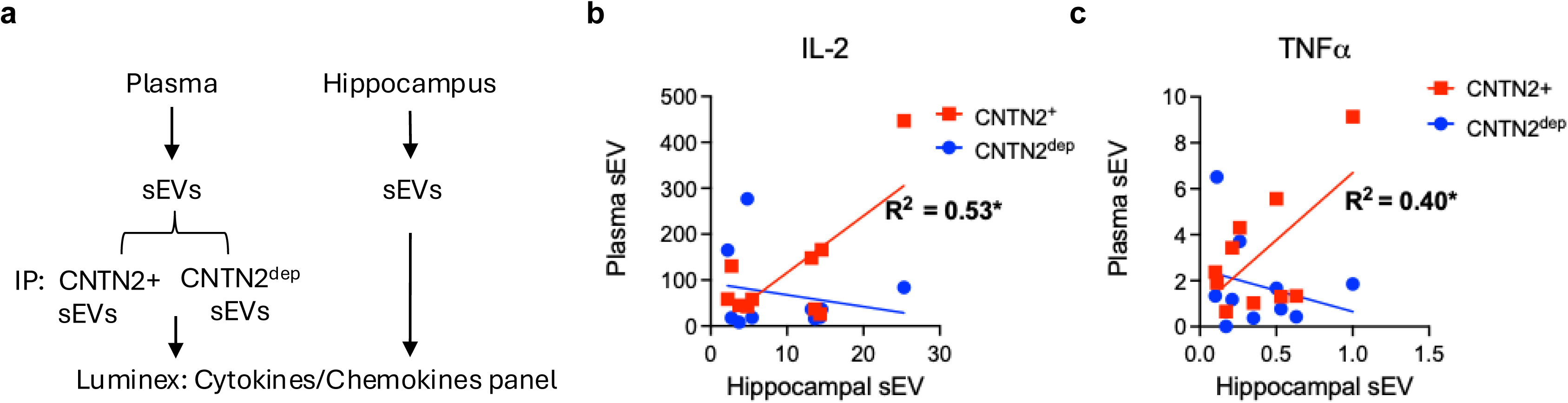
Validation of correlations between plasma BDEVs and brain small EVs (sEVs) in postnatal day 7 rats. (**a**) Schematic illustrating the extraction and analysis of exosomes from plasma and hippocampus. (**b-c**) TNFα and IL-2 showed positive correlations for CNTN2+ (red), but not CNTN2^dep^ (blue) BDEVs extracted from plasma. Linear regression analysis, with r coefficient and p value shown for the CNTN2+ data, n=10.

### Maternal and newborn demographics

Based on the Institute of Medicine (IOM) and American College of Obstetrician and Gynecologists (ACOG) clinical guidelines (Table 1a) (35), group numbers and demographics by pre-pregnancy weight and gestational weight gain (GWG) classifications are detailed in Tables 1b and 1c, respectively. Pre- and post-pregnancy weight, pre-pregnancy BMI and maternal age were significantly different between the two groups with OWO participants being heavier and younger than non-obese controls (Table 1b). While there were notable individual differences with some OWO participants having little to no weight gain, no overall significant difference in gestational weight gain (GWG) during pregnancy was observed between control and OWO groups (Tables 1b-c).

### Characteristics of human plasma BDEVs

While BDEV showed bell-shaped distribution curves ranging between 40 to 200 nm in diameters, the average peak diameter was 80.7 nm in the control group and 92.0 nm in the OWO group. On average, 90.7% of particles in the control group and 84.8% in the OWO group were distributed within their respective peak ranges, consistent with small EV classification (14, 38). The OWO had larger particle diameter than the control group (Supplemental Table 2, Supplemental Figure 2). There was no group difference in BDEV concentrations or proportion of particle distributions.

### Correlations among plasma and BDEV analytes

Umbilical cord plasma and BDEV-associated cytokines, chemokines, nerve growth factors, iron regulators, and neural cell functional markers were quantified and revealed differences between groups in both compartments. Compared to non-obese controls, plasma samples from OWO participants had elevated CRP (Table 2, plasma), consistent with a total body non-infectious inflammatory condition. Within BDEVs, more quantifiable analytes showed significant differences between groups, indicating that BDEVs might provide a better representation of brain-related changes compared to plasma. Changes include higher BDNF, CX3CL1, ferritin, hepcidin, and PARK7 accompanied by lower CCL11, CRP, TNFα, TfR, and S100B in the OWO group compared to the control group (Table 2).

**Table 2.** Concentrations of analytes from plasma and brain-derived extracellular vesicles from cord blood.

| Source | Analyte | Control<br>Mean ± SD (n) | OWO<br>Mean ± SD (n) | *t-test<br>P-value | *Welch's t-test<br>or<br>Mann-Whitney U-test.<br>Significant P-values<br>are in bold. |
| --- | --- | --- | --- | --- | --- |
| Plasma | BDNF (ng/mL) | 0.48 ± 0.45 (19) | 0.28 ± 0.15 (23) | 0.34 | |
| | CCL11 (Eotaxin) (ng/mL) | 0.39 ± 0.13 (19) | 0.33 ± 0.06 (32) | 0.11 | |
| | CRP (ng/mL) | 9.89 ± 6.30 (18) | 16.6 ± 10.9 (35) | **0.04** | |
| | CX3CL1 (ng/mL) | 7.99 ± 3.88 (17) | 9.21 ± 5.13 (28) | 0.37 | |
| | Ferritin (ng/mL) | 57.3 ± 36.4 (20) | 41.0 ± 26.1 (34) | 0.12 | |
| | Hepcidin (ng/mL) | 13.5 ± 9.7 (20) | 9.35 ± 3.93 (20) | 0.49 | |
| | PARK7 (ng/mL) | 13.1 ± 7.0 (17) | 10.3 ± 5.50 (23) | 0.17 | |
| | S100B (ng/mL) | 2.28 ± 1.25 (19) | 1.69 ± 0.52 (23) | 0.24 | |
| | TfR (μg/mL) | 16.2 ± 9.32 (21) | 16.0 ± 8.30 (34) | 0.94 | |
| | TNF-alpha (pg/mL) | 8.41 ± 4.32 (19) | 7.83 ± 4.63 (23) | 0.67 | |
| BDEs | BDNF (pg/10<sup>9</sup> particles) | 1.23 ± 0.09 (19) | 2.72 ± 2.07 (34) | **<0.001** | |
| | CCL11 (Eotaxin) (pg/10<sup>9</sup> particles) | 114.0 ± 54.8 (21) | 66.6 ± 51.5 (36) | **0.003** | |
| | CRP (pg/10<sup>9</sup> particles) | 41.0 ± 25.6 (21) | 15.6 ± 11.4 (33) | **<0.001** | |
| | CX3CL1 (ng/10<sup>9</sup> particles) | 0.53 ± 0.28 (20) | 0.89 ± 0.52 (33) | **0.006** | |
| | Ferritin (pg/10<sup>9</sup> particles) | 49.8 ± 39.6 (21) | 103 ± 40.1(36) | **<0.001** | |
| | Hepcidin (pg/10<sup>9</sup> particles) | 66.3 ± 36.2 (19) | 105 ± 62.0 (35) | **0.01** | |
| | PARK7 (pg/10<sup>9</sup> particles) | 276 ± 190 (21) | 413 ± 185 (36) | **0.01** | |
| | S100B (pg/10<sup>9</sup> particles) | 144 ± 58 (20) | 87.0 ± 29.2 (34) | **<0.001** | |
| | TfR (ng/10<sup>9</sup> particles) | 2.76 ± 1.25 (21) | 1.69 ± 0.95 (36) | **<0.001** | |
| | TNF-alpha (pg/10<sup>9</sup> particles) | 1.49 ± 0.87 (20) | 0.42 ± 0.36 (34) | **<0.001** | |

Spearman r correlation analyses revealed distinct patterns of associations among analytes between groups as well as compartments (Fig. 3). In the control group, iron markers (ferritin, TfR, hepcidin) showed more robust associations with inflammatory markers (TNFα, CRP, CCL11, and CX3CL1) in BDEVs than plasma, indicating higher responsivities of these markers to inflammatory state in the brain compartment (Figs. 3a,b). In the OWO group, iron and inflammatory markers showed positive associations in the plasma compartment (Fig. 3c), and their associations in the brain compartment were also enhanced (Fig. 3d). Compared to the control group, the OWO group showed changes in pattern of associations among markers for both inflammation and iron regulation (Figs. 3c-d vs a-b). Notably, no significant correlation was observed between the plasma and the BDEV compartments (Supplemental Figure 3).

**Fig. 3:**
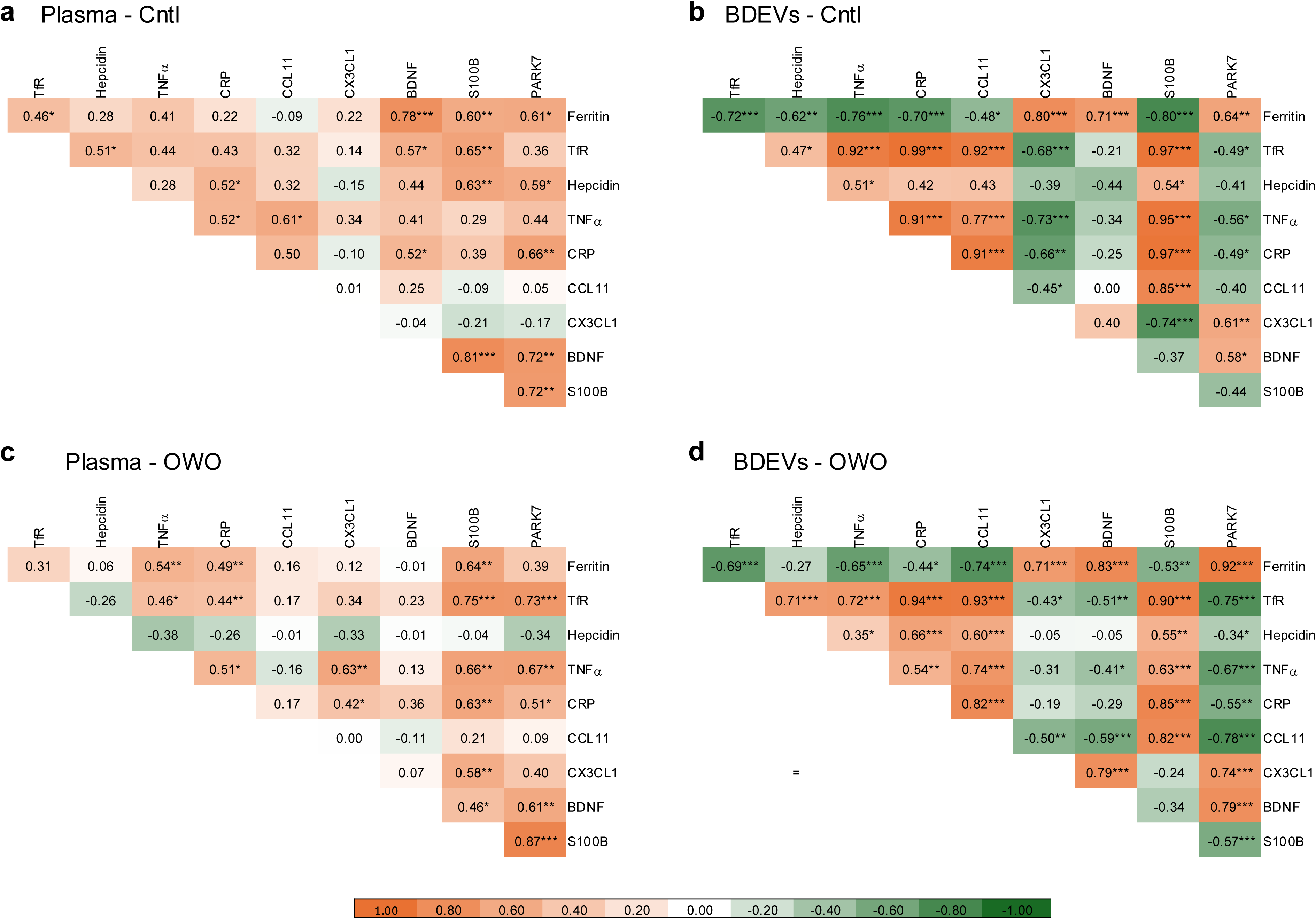
Associative analyses of quantified analytes in cord blood plasma and brain-derived extracellular vesicles. Spearman r correlation matrices showed significant associations among analytes in both control (a) and OWO (b) groups. BDEVs showed more robust and expected associations among iron markers such as inverse relationships of ferritin with TfR and hepcidin as well as among pro-inflammatory markers (e.g., TNFa, CRP, CCL11). Associative changes among the analytes between OWO and control groups uncover the effects of maternal obesity on the newborn’s physiological homeostasis. Asterisk denotes *p<0.05, **p<0.01, ***p<0.001.

To further evaluate the OWO-induced changes of systemic iron accretion and inter-organ iron trafficking in the offspring, regression analyses were performed to assess the associations between ferritin (indicative of iron preservation or sequestration) and TfR (indicative of cellular iron demand) (39) in the periphery and CNS. While the frequency distribution of ferritin and TfR values was not different between control and OWO group in plasma, they shifted to higher and lower concentrations, respectively, in the OWO compared to the control group in BDEVs (Figs. 4a-d). Using plasma ferritin as an independent variable to evaluate their responses in brain, regression curves showed different responses between the two groups for BDEV ferritin and TfR (Figs. 4e-f). In the control group, BDEV ferritin increased, and BDEV TfR decreased, as plasma ferritin decreased (Figs. 4e-f, green lines). In contrast, the OWO group showed an apparent drop in BDEV ferritin as plasma ferritin declined below 50 ng/mL, approximately the inflection point at which brain iron becomes compromised (40, 41) (Figs. 4e-f, orange lines).

**Fig. 4:**
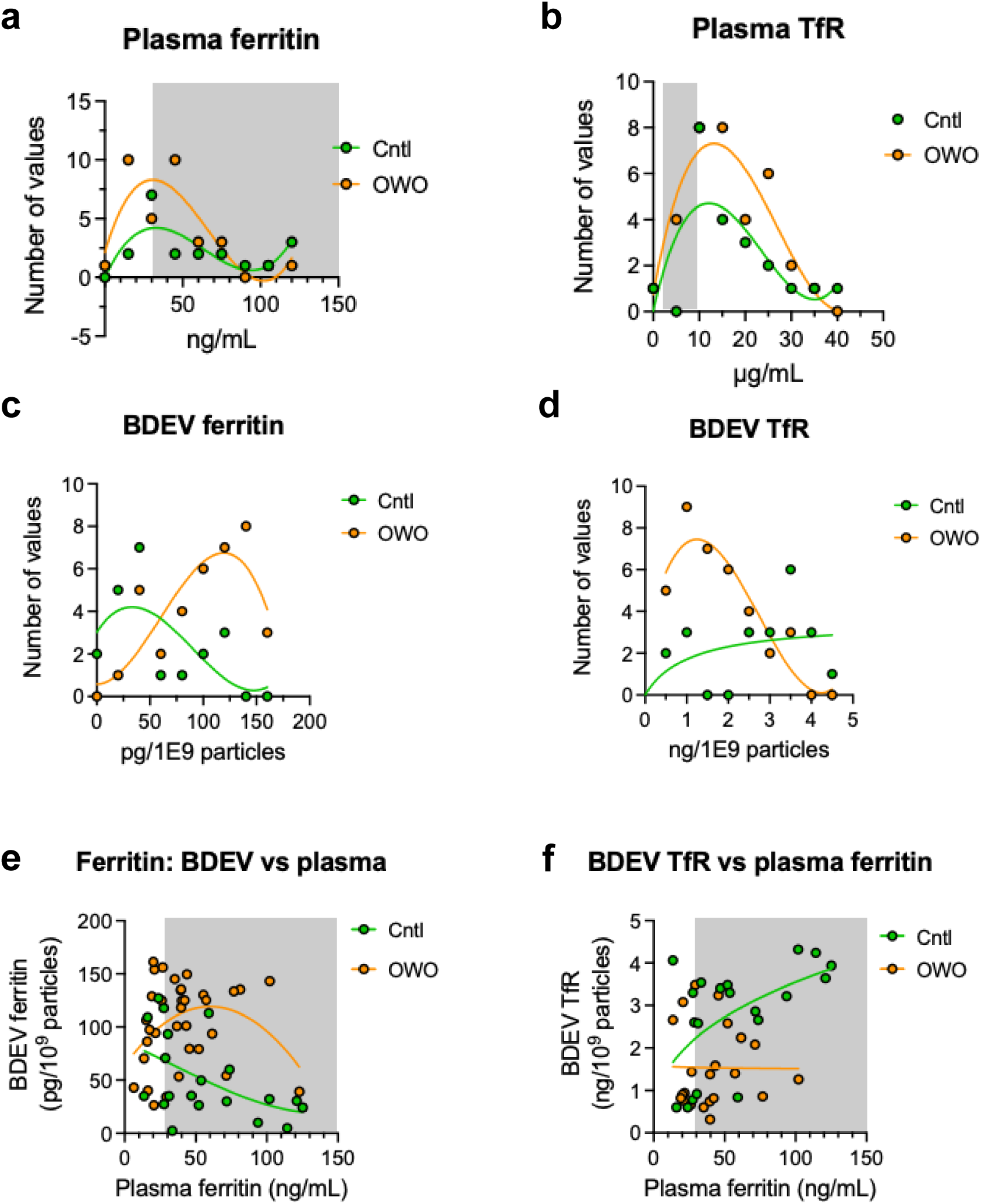
Associations between systemic and CNS iron accumulation and trafficking. (**a-d**) Frequency distributions of ferritin and TfR concentrations in plasma and BDEVs. Compared to control (Cntl), overweight-obese (OWO) group cord plasma and BDEVs showed a shift to higher ferritin and lower TfR concentrations, respectively, suggesting increased iron storage and decrease iron demand in the brain. Grey shades represent normal physiological ranges of plasma ferritin and TfR. No established normal range of ferritin and TfR in the brain. (**e-f**) Regression analyses with plasma ferritin as the independent variable. Results showed difference responses of BDEV ferritin and TfR between control and OWO groups. Non-linear regression with post hoc extra sum-of-squares F test for curve fitness comparisons between groups. Separate lines indicate significant difference between curves (p<0.05).

### Anti-inflammatory and neuroprotective signatures are upregulated in BDEVs of cord bloods from OWO pregnancies

To evaluate the maternal obesity-induced changes in brain inflammatory signatures in the newborns, concentrations of cytokines and chemokines in cord plasma BDEVs were assayed. The OWO group showed an elevated CX3CL1 concentration, but reduced concentration of TNFα, CRP, and CCL11 compared to control group (Fig. 5a). Given the known role of neuronal-derived CX3CL1 in inhibiting microglial TNFα expression (42, 43), BDEV CX3CL1 was used as independent variable to evaluate the response of BDEV TNFα. TNFα showed a negative association with CX3CL1 in the control group; however, this association was significantly diminished in the OWO group (Fig. 5c). Consistent with known positive associations of TNFα with CRP and CCL11 in the CNS (44–46), non-linear regression analyses BDEV data using TNFα as the independent factor showed direct relationships with CRP and CCL11 (Figs.5d-e). There was no difference between OWO and control groups for the associations between CRP, CCL11, and TNFα, indicating by shared responsive curves (Figs.5d-e).

**Fig. 5:**
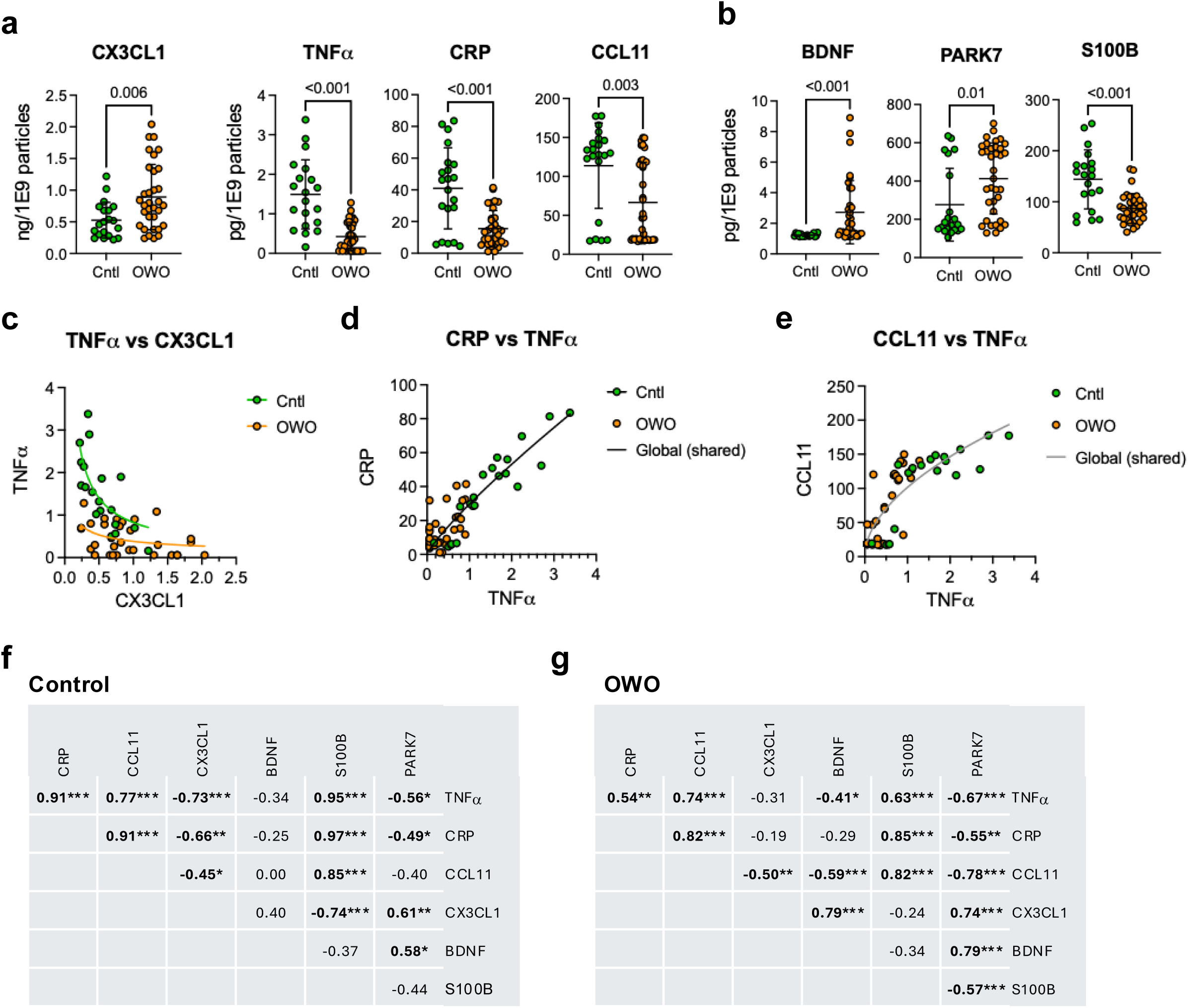
Changes of cytokines, chemokines, and downstream factors in cord plasma BDEVs extracted immediately post birth. (**a**) BDEV cytokines and chemokines of newborns from overweight-obese (OWO, orange dots) group showed higher neuronal-specific anti-inflammatory CX3CL1 and lower pro-inflammatory TNFα, CRP and CCL11 compared to newborns of non-obese control (Cntl, green dots) group. (**b**) Comparisons of BDEV markers for neural cell functions showed higher concentration of BDEV BDNF and PARK7, and lower S100B concentration in OWO compared to Cntl group. Values are Mean ± SD, Welch’s t-test or Mann-Whitney U-test. (**c**) BDEV CX3CL1 and TNFα showed an inverse association with a lower responsivity of TNFα in the OWO group. (**d-e**) CRP and CCL11 showed positive associations with TNFα. There was no difference between OWO and control groups. Non-linear regression with post hoc extra sum-of-squares F test for curve fitness comparisons between groups. (**f-g**) Spearman’s r matrices showed associations between inflammatory and cell function markers in control and OWO groups. Bold typeface denotes significant association. Asterisk denotes: *p<0.05, **p<0.01, ***p<0.001.

To further assess downstream effects of the altered levels of BDEV cytokines and chemokines, concentrations of markers for neuroprotection and neuroplasticity (BDNF (47, 48)), metabolite-mediated cell signaling and anti-oxidative stress (PARK7 (49, 50)), and astrogliosis (S100B (51, 52)) were analyzed in cord plasma BDEVs. Compared to controls, OWO BDEVs had higher BDNF, higher PARK7, and lower S100B concentrations (Fig. 5b). To determine the association among the expression changes of BDEV cytokines/chemokines and downstream factors (BDNF, PARK7, S100B), Spearman’s r correlations were performed. Compared to the control group, OWO group showed increased strengths of inverse associations between BDEV BDNF and PARK7 with TNFα and CCL11, as well as direct associations with CX3CL1 (Figs. 5f-g). BDEV S100B showed negative associations with TNFα, CRP and CCL11 and positive associations with CX3CL1 and PARK7 (Figs. 5f-g).

### Overweight and obese pregnancy alters inflammatory signatures in umbilical cord plasma

Given prior evidence of elevated pro-inflammatory signatures in newborns of mothers with obesity (53, 54), peripheral immune response was also assessed in this studied cohort. Plasma expression level of inflammatory cytokines (TNFα, CRP) and chemokines (CCL11, CX3CL1) were quantified (Figs. 6a-b). Results showed a higher plasma CRP concentration in the OWO compared to the control group (Fig. 6a). To assess the association between plasma CRP with TNFα and CX3CL1, regression (non-linear) analyses were performed, which showed a direct relationship between plasma CRP and TNFα with a more robust response, indicative of a higher sensitivity, in the OWO group compared to the control group (Fig. 6c). There was no association between plasma CRP and CX3CL1 (Fig. 6d).

**Fig. 6:**
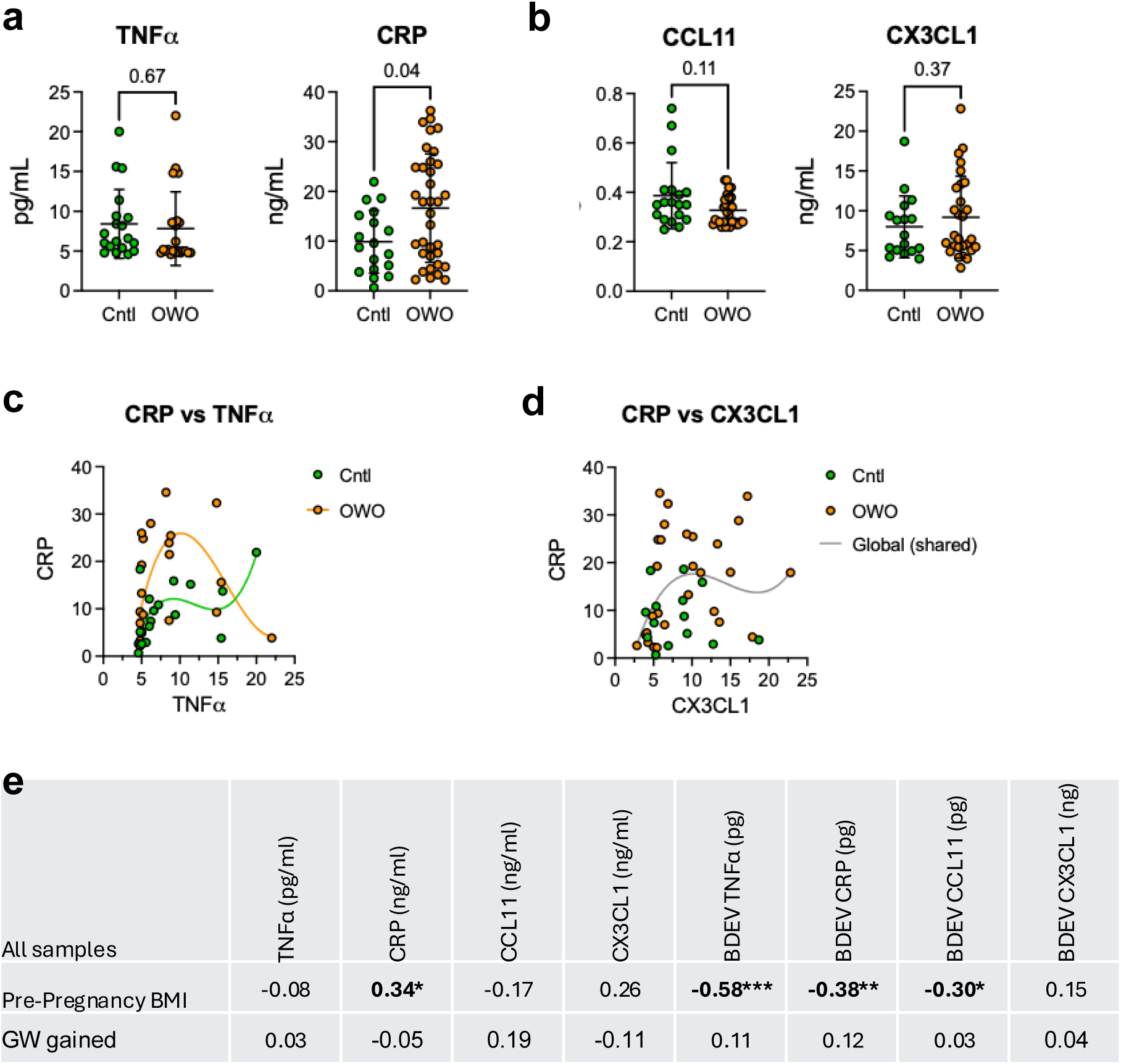
Elevated CRP in cord plasma of the OWO group. (**a**) Plasma concentrations of pro-inflammatory cytokines. OWO group had a higher CRP level than control (Cntl) group. (**b**) No change in level of anti-inflammatory chemokine CX3CL1 between groups. (**c-d**) Using TNFα or CX3CL1 as independent variables to evaluate responses of CRP, the OWO group showed a more robust CRP response than control group to TNFα. There was no association between CRP and CX3CL1. Non-linear regression with post hoc Extra Sum-of-Squares F test for curve fitness comparisons between groups. Separate lines indicate significant difference between curves (p<0.05).

To determine the association between pre-pregnancy BMI and systemic plasma and CNS cytokines/chemokines, Spearman’s correlations were performed. While pre-pregnancy BMI is positively associated with cord plasma CRP level, it is negatively associated with BDEV TNFa, CRP, and CCL11 (Fig. 6e). There was no association between gestational weight gained during pregnancy and these cytokines/chemokines.

### Elevated brain iron storage and reduced neuronal iron uptake in newborns of mothers with obesity

To evaluate the effects of maternal OWO on brain iron regulation in newborns, changes in BDEV ferritin, TfR, and hepcidin were analyzed between control and OWO mothers. OWO BDEVs had higher levels of ferritin and hepcidin, and lower levels of TfR compared to controls (Figs. 7a-c). Using BDEV hepcidin as an independent variable to evaluate the responses of BDEV ferritin and TfR, regression (non-linear) analyses revealed different responses between control and OWO groups. In the control group, BDEV ferritin and TfR declined and rose sharply, respectively, and then plateaued as BDEV hepcidin increased to about 50 pg/1E^9^ particles (Figs. 7d-e, green lines). In contrast, the OWO group showed a gradual decrease and increase of ferritin and TfR, respectively, as hepcidin increased (Figs. 7d-e, orange lines), indicating a lower sensitivity of ferritin and TfR to hepcidin in the OWO group. While the inverse correlations were found in both groups for BDEV ferritin and BDEV TfR, the OWO group showed a significantly less negative slope than that of the control group (Fig. 7f, differences between slopes). Pre-pregnancy BMI, but not gestational weight gained (GWG) during pregnancy, showed positive and negative associations with BDEV ferritin and TfR (Fig. 7g).

**Fig. 7:**
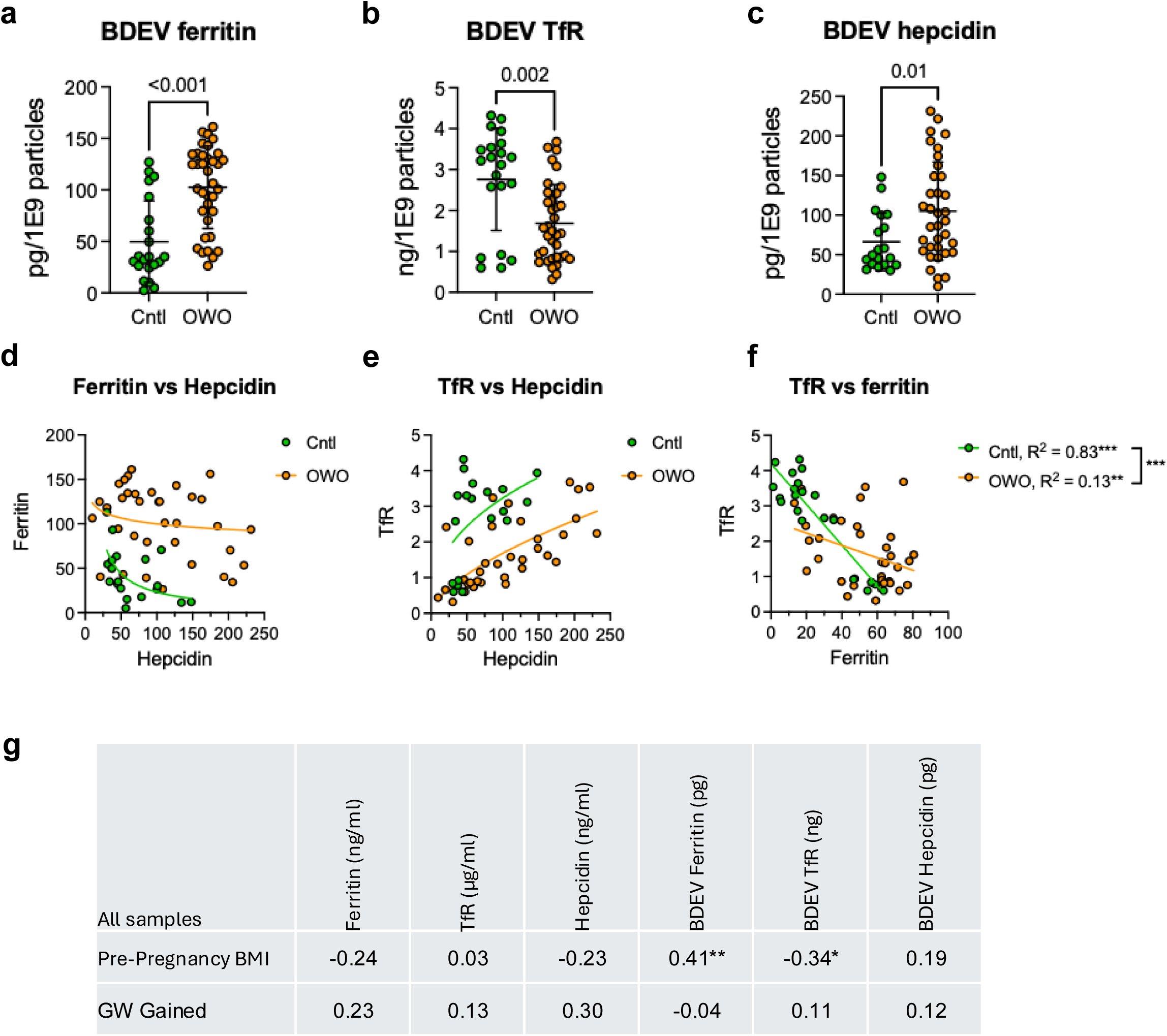
Differential brain iron responses between control and OWO groups. (**a-c**) Compared to control (Cntl) group, OWO group showed higher concentrations of ferritin and hepcidin, and lower concentration of TfR in BDEVs. Values are Mean ± SD, Welsch’s *t-*test or Mann-Whitney U-test. (**d-e**) Regression analyses using hepcidin as the independent variable showed differential responses of ferritin and TfR between control and OWO groups. Non-linear regression with post hoc Extra Sum-of-Squares F test for curve fitness comparisons between groups. Separate lines indicate significant difference between curves (p<0.05). (**f**) TfR and ferritin showed an inverse correlation with different (linear) regression curves. Asterisks denote: **p<0.01, ***p<0.001.

## Discussion

In this study, we demonstrated the utility of plasma BDEVs as a non-invasive approach for assessing CNS physiology using both preclinical and human umbilical cord blood samples. We further used this approach to evaluate NINI and nID in newborns of women with OWO. The preclinical data indicated that plasma BDEVs are derivable from brain tissue (e.g., hippocampus) and that the contents in plasma BDEVs can accurately reflect brain status. The clinical data indicated that analytes related to ID and inflammation showed stronger correlations in BDEVs than in plasma, suggesting that markers carried by BDEVs can represent brain physiology more accurately than those in plasma. Employing this approach, we found that otherwise healthy newborns delivered by C-section from uncomplicated pregnancies of overweight-obese (OWO) women had elevated levels of BDEV anti-inflammatory chemokine CX3CL1 and neuroprotective factor (BDNF, PARK7), and reduced levels of BDEV pro-inflammatory TNFα, CRP, and Eotaxin (CCL11) and an astrogliosis marker S100B. These differences were accompanied by elevated levels of BDEV signal for iron sequestration (i.e., hepcidin and ferritin) along with reduced signal for neuronal iron accretion (i.e., TfR). BDEV ferritin and BDEV TfR concentrations were inversely and directly related to BDEV proinflammatory markers (TNFα, CRP, Eotaxin), respectively, and as expected inversely related to each other. Given the previously reported pro-inflammatory state in OWO pregnancies (25, 53, 54), which is further supported in this study with the finding of higher plasma CRP levels in the cord blood of OWO compared to non-obese controls, we postulate that these BDEV changes reflect a protective response by the fetal brain to the peripheral pro-inflammatory state.

The discovery of quantifiable BDEV biomarkers to index fetal NINI and nID in a non-invasive sample offers a new set of tools to evaluate brain status in newborn infants at risk for long-term adverse neurodevelopmental outcomes from a diverse group of *in utero* conditions (e.g., preeclampsia, chorioamnionitis, nutritional deficiencies, teratogen exposures). BDEV contents can be measured during critical window of peripartum neurodevelopment when postnatal interventions could have a greater impact. The data further suggest that peripheral blood biomarkers may be inaccurate surrogates for changes in the brain, most likely due to dilutional contributions from other organs to the plasma pool, and that assumptions of neurodevelopmental risk based on peripheral biomarkers of NINI or nID should be interpreted with caution.

The prevalence of gestational obesity, defined as either high pre-pregnancy BMI, excessive gestational weight gain or both, rose from +20% in 1990s to +40% in 2019 (55), raising concerns for long-term health risks for the offspring (56). However, mechanisms underlying these health risks remain understudied, particularly regarding the link between maternal obesity and neurodevelopmental disorders in the offspring (57, 58). The key limitation of these studies is that the effects of maternal obesity on offspring brain are inferred from biomarkers measured in the peripheral circulation, which can be unreliable, thus obscuring brain specific vulnerability or protection. This study leveraged BDEVs, nanoparticles secreted and trafficked from the CNS that were purified from cord blood, to evaluate obesity-mediated differences in molecular markers (i.e., cytokines, chemokines, nerve growth factors, and indices of iron status) of relevant pathophysiologic conditions. Our findings uncover differences between maternal obesity and offspring peripheral vs brain physiological states than previously suspected based only on peripheral data. Consistent with prior findings (53), newborns of OWO mothers have elevated plasma CRP, a standard biomarker of pro-inflammation in the periphery; however, the brain-specific data provide evidence for active anti-inflammatory processes in microglia, reduction of astrogliosis, and promotion of neuroprotection/neuroplasticity. In addition, the newborn brain of OWO mothers has evidence of a functional nID evidenced by enhanced ferritin iron storage and reduced neuronal iron import via TfR (59). These findings provide insight about potential mechanisms of action among neural cell types in the developing brain of newborns of obese mothers. Pre-pregnancy BMI, but not GWG during pregnancy, is likely a contributor to functional neuronal ID and immune responses in the OWO offspring, suggesting that pre-existing adiposity may set the stage for a pro-inflammatory milieu during gestation that impairs iron regulation and activates anti-inflammatory responses in the developing fetal brain.

The anti-inflammatory response of the fetal/newborn brain was a novel finding that illustrates the disconnect between peripheral and centrally derived biomarkers. CX3CL1/CX3CR1 signaling represents a neuron-microglia crosstalk, where the neuronal specific chemokine CX3CL1 activates its microglial-specific receptor CX3CR1 to inhibit TNFα expression, promoting anti-inflammatory and neuroprotective processes (43, 60, 61). The inverse correlation between CX3CL1 and TNFα in BDEVs is consistent with the known inhibition of TNFα by CX3CL1 signaling. This correlation was less robust and non-significant in cord plasma BDEVs of the OWO group, indicating a decrease in sensitivity of fetal CX3CL1 signaling. This reduced sensitivity was also observed for BDEV S100B, which indexes astrocyte activity and regulates microglial TNFα (52, 62). The data showed that OWO offspring have a different CNS immune responsiveness compared to offspring of non-obese controls. Whether this is a compensatory adaptive response or evidence of dysregulation is unclear and requires further investigation. These potential protective molecular responses in the brain are further supported by elevated concentrations of an anti-oxidative stress marker, PARK7 (49, 50) and neuroprotective/ neuroplasticity marker, BDNF (47), while reduced adverse astrocyte activity (astrogliosis) as indicated by lower S100B levels (51). While the link between maternal obesity and the programming of peripheral inflammation in offspring has been documented (53, 63), this study provides the first documentation in human newborns that a reprogramming occurs in the brain as well.

Maternal obesity is a known risk factor for peripheral ID in the offspring (64); however, there is limited ability to study multiple burden of risk, including NINI and nID and associated neurodevelopmental impairments in newborns (64). That is because, until this point, there have not been reliable non-invasive markers to parse out these molecular mechanisms in asymptomatic infants. Our findings of reduced TfR concomitant with elevated ferritin and hepcidin in BDEVs provide evidence that BDEVs can index brain iron regulation and that functional ID can be detected in the CNS of OWO offspring, similar to a biological process known as anemia of inflammation (high hepcidin, high ferritin, low TfR (65, 66)). We propose that these effects were driven by elevated levels of BDEV hepcidin in the OWO-exposed fetal brain. Given that CX3CL1/CX3CR1 signaling has been hypothesized to promote hepcidin synthesis in microglia and has been associated with subsequent iron accumulation in neurons (67), our observed positive correlations between BDEV hepcidin with BDEV TNFα, CRP, and CCL11 (Figs. 3d,e) only in the OWO offspring provide additional evidence supporting this hypothesis. Taken together, our findings suggest that regulation of iron homeostasis in the CNS relies on the local environment (high BDEV CX3CL1, high BDEV hepcidin), which itself is amenable to the peripheral regulation (Fig. 4). These findings provide evidence for the presence of brain functional ID preceding the signatures of peripheral functional ID, which is consistent with the principle that iron is prioritized first to the production of the red cells at the expense of brain cells (40, 68).

Strengths of this study include the large number of samples, the highly sensitive assays, and the use of both plasma and BDEV chemokines and iron proteins. However, several limitations exist. First, samples were collected from unlabored elective cesarean deliveries and findings may be limited to that group, but this could also be considered a strength due to the alterations of cytokines in clinical chorioamnionitis. Second, the cross-sectional nature of the analysis could not address whether the identified changes were long-term adaptive or dysregulated responses to maternal obesity. Third, the impact of newborn sex on findings was not examined and should be studied in a larger cohort.

In conclusion, this study demonstrates the utility and sensitivity of quantifiable BDEV biomarkers for neuroinflammation and brain iron status, two prevalent gestational conditions world-wide that have been associated with neurodevelopmental disorders in later life. This is a major advance given previous reliance on either peripheral biomarkers that may not reliably index the brain physiological states or on direct invasive procedures such as spinal tap or direct tissue assessment. Our findings suggest a working model for the interaction of brain inflammation and iron status in OWO offspring, where altered BDEV molecular markers indicate neuronal functional iron deficiency accompanied by reduced neuroinflammation (Fig. 8). The data provide groundwork for future studies to establish BDEVs as clinical tools in identifying infants at risk for long-term adverse neurodevelopmental outcomes before the development of clinical signs later in life. Such studies could either measure BDEV contents directly or by relating them to more commonly assayed peripheral biomarkers of inflammation or iron status.

**Fig. 8:**
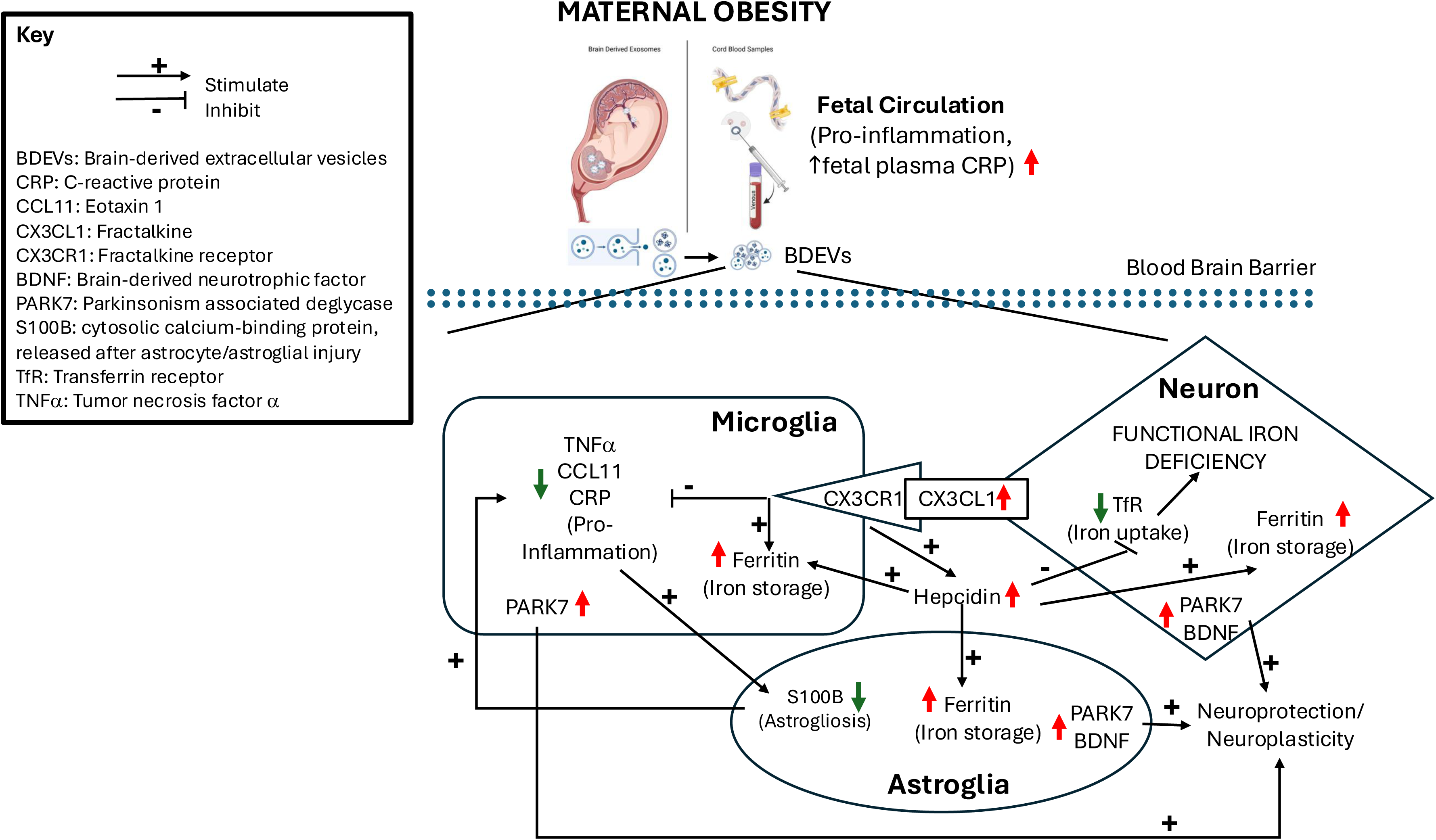
A proposed model of altered brain iron indices and immune responses in fetus/newborn of mother with obesity through altered maternal-fetal metabolic milieu on microglia, astrocytes, and neurons. Maternal obesity can induce the cross-talks among fetal neural cells, including activation of the neuron-microglia CX3CL1/CX3CR1 signaling and astrocyte-microglia S100B signaling, leading to anti-inflammatory responses, including reduction of pro-inflammation (TNFa, CRP, CCL11) and promotion of neuroprotection (BDNF, PARK7) and cell metabolism (PARK7). Conversely, maternal obesity can induce neuronal functional iron deficiency (high ferritin, low TfR). Hepcidin, an acute phase protein and also the master iron regulator may play a role.

## Author contributions

PVT, PJK, and MKG developed the concept and designed the study; SXL, ACH, and PM generated preclinical biospecimens; NJ and PJK provided human biospecimens for the initial pilot assays (data not shown); MLB and SK provided human biospecimens of the studied cohort; MM and ZM conducted the experiments; PVT and MLB analyzed the data; PVT and MLB has directly accessed and verified the underlying data reported in the manuscript; PVT drafted the manuscript; PVT, SXL, ACH, NJ, PJK, MKG, and MLB reviewed and edited the manuscript and assisted with figures; all authors reviewed and approved the manuscript.

This work has not been presented before, but preliminary data were generated after improving methodology from a smaller comparable dataset of otherwise healthy scheduled cesarean deliveries from the Meriter Hospital were presented at the both the 2023 Midwest Society for Pediatric Research and 2024 Pediatric Academic Societies. Dolan N, Tran P, Kling PJ, Georgieff M, Mukhtarova N. Biomarkers of Brain Exosomes from Plasma Displayed a Link between Neonatal Systemic and Central Nervous System Inflammation 10/2023. Tran PV, Dolan N, Mukhtarova N, Kling PJ, Georgieff MK. Biomarkers of Brain Exosomes from Plasma Displayed a Link between Neonatal Systemic and Central Nervous System Inflammation, Pediatric Academic Societies, 2024.

## Acknowledgements

This work was supported in part by the NIH Grant to MKG and PVT (P01-046925), financial support from the Masonic Institute for Developing Brain to PVT and ACH. Financial support from the Hennepin Healthcare Research Institute to ACH. PJK and NJ were supported by the UW Department of Pediatrics, UnityPoint Health Meriter Foundation, and Gerber Foundation. MLB was supported by NIH/NIGMS P20GM103620, NIH/NHLBI R01 HL160980, NIH/NIGMS P30 GM154633. SK has an AAP grant that was used to support the collection of cord blood samples to measure leptin/adiponectin cytokines: American Academy of Pediatrics Resident Research Grant.

We thank Sharon Blohowiak, BS and Natalie Dosch, MD for making available biospecimens used in the initial pilot/feasibility study. We thank Dr. James Marti in Nanotechnology Lab for assisting in the use of NTA instruments. We thank Ms. Fang Zhou for imaging of extracellular vesicles by Transmission Electron Microscopy in the Characterization Facility, University of Minnesota, which receives support in part from the NSF through the MRSEC (Award Number DMR-2011401) and the NNCI (Award Number ECCS-20245124) programs. We thank Michael Ehrhardt and Dr. Angela Mortari-Panoskaltsis of the Peds Cytokine Lab for valuable advice on assays for cytokines and chemokines. We thank our enrolled patients, clinical coordinators (Hannah Kienast, RN and Emily Andersen, RN) for their consented, enrolled, and assisted in the sample collection process.

