## Supplemental Fig.1 for "Utility of brain-derived extracellular vesicles from human umbilical cord blood to measure non-infectious neuroinflammation and functional iron deficiency"

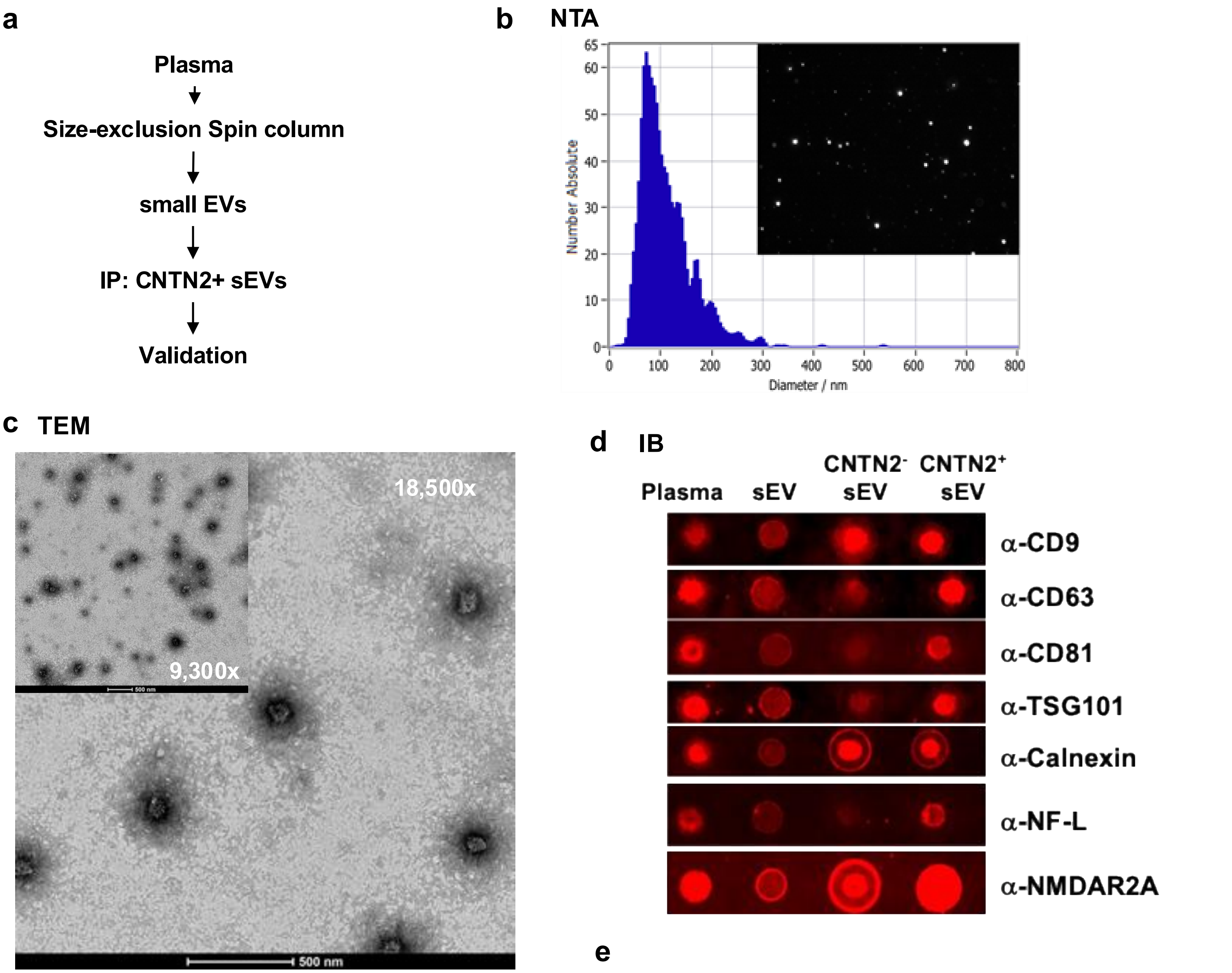


**e**

| Sample | CD9 | CD63 | CD81 | Syn-1 | Flot-1 | TSG101 | Ago-2 | CALR | GAPDH |
|---|---|---|---|---|---|---|---|---|---|
| BDEV1 | 0.40 | 712 | 0.11 | 5.48 | 139 | 2537 | 1474 | 118 | 96.4 |
| BDEV2 | 0.01 | 161 | 0.01 | 6.10 | 0.98 | 475 | 43.5 | 9.47 | 229 |
| BDEV3 | 0.01 | 90.5 | 0.001 | 9.58 | 2.33 | - | - | 11.2 | 67.4 |

Supplemental Figure 1: Isolation, enrichment, imaging, and quantitation of brain-derived small extracellular vesicles (BDEVs) from plasma. (**a**) A Schematic illustration of the method to isolate BDEVs using size exclusion column chromatography and immunoprecipitation (IP) with Contactin-2 (CNTN2). (**b**) BDEVs were qualified with validation for size and concentration by a Nanoparticle Tracking Analyzer (NTA, ZetaView). Representative image shows peak and distribution of BDEV diameters across a range of 40 nm to 250 nm. (**c**) Images of BDEVs captured by transmission electron microscopy (TEM) at 18,500 magnifications. Inset is a TEM image at 9,300 magnifications showing a field of BDEVs at various sizes. (**d**), Expression of appropriate markers for BDEVs, including tetraspanins (CD9, CD63, CD81), biogenesis (TSG101), and neural cells (CNTN2, NF-L). (**e**) Confirmation of exosome characteristics using Milliplex Human Exosome Characterization panel (HEXSM-170-PMX) and Luminex platform. The presence of Syn-1 and Flot-1 indicates that particles underwent cargo sorting process during biogenesis, including incorporation of Calreticulin (CALR) and GAPDH.
