## Supplemental Fig.2 for "Utility of brain-derived extracellular vesicles from human umbilical cord blood to measure non-infectious neuroinflammation and functional iron deficiency"

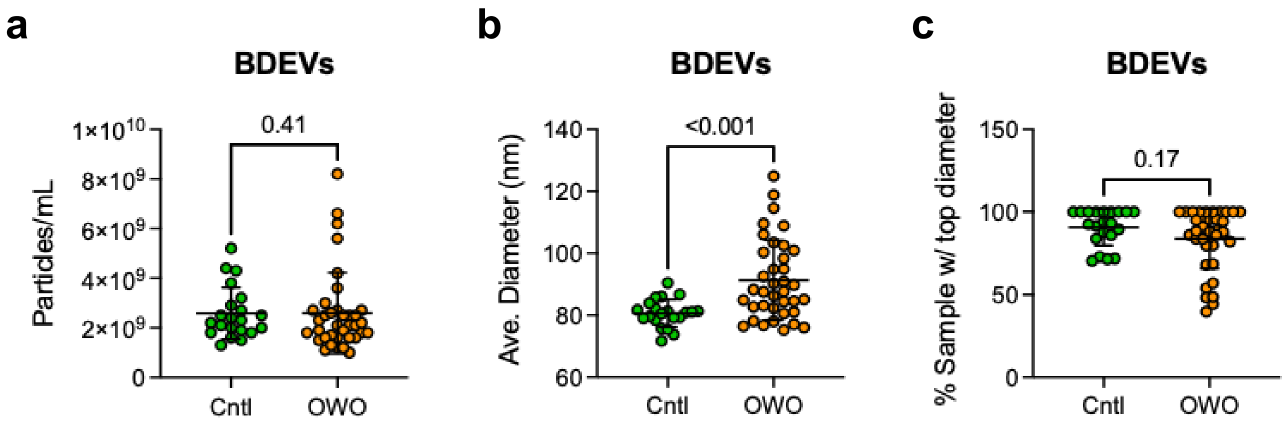


**Supplemental Figure 2** – BDEV characteristics assessed by Nanoparticle Tracking Analyzer. (**a**) Concentrations of BDEVs range between $1.0x10^9$ and $1.0x10^{10}$ particles/mL. (**b**) BDEV sizes from the OWO are larger than those from the Cntl group, indicating potentially larger cargo loads. (**c**) > 80% particles in each BDEV isolates have a peak diameter between 80-100 nm.
