## Supplemental Table 1 for "Utility of brain-derived extracellular vesicles from human umbilical cord blood to measure non-infectious neuroinflammation and functional iron deficiency"

| Target | Application | Host Species | Clonality | Cat. # | Manufacturer |
|---|---|---|---|---|---|
| CNTN2 | IP | Rabbit | Polyclonal | PA5-101541 | Invitrogen |
| CD9 | DB primary | Mouse | Monoclonal | 10626D | Invitrogen |
| CD63 | DB primary | Mouse | Monoclonal | 10628D | Invitrogen |
| CD81 | DB primary | Mouse | Monoclonal | 10630D | Invitrogen |
| TSG101 | DB primary | Mouse | Monoclonal | MA1-23296 | Invitrogen |
| Calnexin | DB primary | Mouse | Monoclonal | MA5-31501 | Invitrogen |
| NF-L | DB primary | Mouse | Monoclonal | Ab273441 | Abcam |
| NMDAR2A | DB primary | Mouse | Monoclonal | MA5-27693 | Invitrogen |

Supplemental Table 1: Information of immunoprecipitation (IP) and dot blot (DB) primary antibodie[illegible] s used in the study.
