## Supplemental Table 2 for "Utility of brain-derived extracellular vesicles from human umbilical cord blood to measure non-infectious neuroinflammation and functional iron deficiency"

**Supplemental Table 2** - Average size and concentration of BDEVs purified from cord plasma

| Control | | | |
|---|---|---|---|
| Sample ID | Concentration (Particles/ml) | Peak Diameter (nm) | % Sample within Peak |
| 74 | 2.00E+09 | 81.3 | 71.8 |
| 93 | 1.30E+09 | 90.4 | 91.6 |
| 95 | 5.20E+09 | 80.9 | 100.0 |
| 103 | 2.30E+09 | 75.7 | 100.0 |
| 105 | 4.30E+09 | 86.0 | 100.0 |
| 123 | 4.40E+09 | 82.1 | 94.0 |
| 139 | 1.50E+09 | 73.8 | 100.0 |
| 143 | 1.80E+09 | 78.4 | 72.8 |
| 149 | 2.20E+09 | 79.6 | 87.4 |
| 159 | 1.80E+09 | 86.7 | 70.4 |
| 186 | 3.20E+09 | 75.4 | 85.9 |
| 192 | 1.60E+09 | 81.7 | 100.0 |
| 203 | 2.90E+09 | 81.3 | 100.0 |
| 211 | 2.50E+09 | 79.0 | 89.6 |
| 220 | 2.50E+09 | 79.3 | 100.0 |
| 226 | 2.00E+09 | 85.7 | 92.6 |
| 236 | 3.80E+09 | 81.1 | 100.0 |
| 241 | 1.90E+09 | 71.7 | 100.0 |
| 259 | 2.10E+09 | 80.5 | 83.8 |
| 264 | 2.70E+09 | 83.9 | 71.4 |
| 273 | 2.40E+09 | 79.2 | 93.3 |
| **Average** | **2.59E+09** | **80.652** | **90.695** |

| Overweight-obese (OWO) | | | |
|---|---|---|---|
| Sample ID | Concentration (particles/mL) | Peak Diameter (nm) | % Sample within Peak |
| 88 | 5.60E+09 | 94.9 | 100.0 |
| 92 | 1.80E+09 | 75.2 | 94.8 |
| 98 | 6.20E+09 | 82.7 | 100.0 |
| 101 | 2.50E+09 | 77.1 | 100.0 |
| 104 | 2.10E+09 | 100.9 | 84.5 |
| 106 | 1.60E+09 | 87.6 | 91.2 |
| 114 | 2.70E+09 | 85.1 | 86.1 |
| 118 | 2.70E+09 | 78.0 | 95.4 |
| 119 | 1.60E+09 | 119.4 | 86.0 |
| 121 | 6.60E+09 | 118.8 | 48.2 |
| 130 | 1.60E+09 | 109.6 | 88.8 |
| 133 | 2.50E+09 | 90.9 | 86.4 |
| 136 | 2.20E+09 | 84.4 | 68.1 |
| 137 | 2.30E+09 | 76.0 | 95.6 |
| 151 | 1.70E+09 | 106.0 | 39.7 |
| 155 | 8.20E+09 | 92.6 | 83.7 |

| | | | |
|---|---|---|---|
| 161 | 1.10E+09 | 100.3 | 100.0 |
| 164 | 1.50E+09 | 76.4 | 56.9 |
| 184 | 1.20E+09 | 98.3 | 100.0 |
| 188 | 1.80E+09 | 82.3 | 88.2 |
| 194 | 3.00E+09 | 86.2 | 100.0 |
| 197 | 2.70E+09 | 81.0 | 100.0 |
| 210 | 4.20E+09 | 102.5 | 100.0 |
| 213 | 2.50E+09 | 76.8 | 79.7 |
| 230 | 2.50E+09 | 78.1 | 94.3 |
| 231 | 2.60E+09 | 87.7 | 82.0 |
| 238 | 1.90E+09 | 124.9 | 44.3 |
| 239 | 1.60E+09 | 83.1 | 88.6 |
| 242 | 3.60E+09 | 94.8 | 88.1 |
| 246 | 1.30E+09 | 84.6 | 68.5 |
| 250 | 2.10E+09 | 80.6 | 100.0 |
| 257 | 1.60E+10 | 103.4 | 100.0 |
| 258 | 1.20E+09 | 88.2 | 48.6 |
| 261 | 1.80E+09 | 84.9 | 95.0 |
| 266 | 1.90E+09 | 90.2 | 83.7 |
| 267 | 1.60E+09 | 89.8 | 86.1 |
| **Average** | **3.00E+09** | **90.925** | **84.792** |
